# Genomic Analyses of the Metastasis-derived LNCaP, VCaP, and PC3-AR Prostate Cancer Cell Lines

**DOI:** 10.1101/2021.06.25.449904

**Authors:** Karolina Sienkiewicz, Chunsong Yang, Bryce M. Paschal, Aakrosh Ratan

**Author notes:** Correspondence should be addressed to B.M.P. and A.R.

## Abstract

The lymph node metastasis-derived LNCaP, the bone metastasis-derived PC3 (skull), and VCaP (vertebral) cell lines are widely used preclinical models of human prostate cancer (CaP) and have been described in >19,000 publications. Here, we report on short-read whole-genome sequencing and genomic analyses of LNCaP, VCaP, and PC3 cells stably transduced with WT AR (PC3-AR). LNCaP cells are composed of multiple subpopulations, which results in non-integral copy number states and a high mutational load when the data is analyzed in bulk. All three cell lines contain pathogenic mutations and homozygous deletions in genes involved in DNA mismatch repair, along with deleterious mutations in cellcycle, Wnt signaling, and other cellular processes. Furthermore, LNCaP cells contain a missense mutation in a well-known CaP hotspot of TP53, whereas both PC3-AR and VCaP have truncating mutations in TP53 and do not express p53 protein. In addition, we detect the signatures of chromothripsis of the q arms of chromosome 5 in both PC3-AR and VCaP cells, strengthening the association of TP53 inactivation with chromothripsis reported in other systems. Our work provides a resource for genetic, genomic, and biological studies employing these commonly-used prostate cancer cell lines.

## Introduction

Cancer lines, such as the human-derived LNCaP, VCaP, and PC3 cells are critical tools for prostate cancer (CaP) research. The androgen-dependent LNCaP and VCaP cell lines are derived from a lymph node metastasis and a vertebral metastasis, respectively [1, 2]. The androgen-independent PC3 cell line is derived from a skull metastasis [3]. PC3 cells are AR-negative, but several groups including ours have reintroduced WT androgen receptor (AR) into these cells to study androgen signaling [4–6]. Thus, stable transduction of AR into PC3 cells has been used generate “PC3-AR” cells that express full-length AR protein at levels comparable to endogenous AR in VCaP and LNCaP. Transcriptomic analysis of PC3-AR cells revealed the presence of a large number of 1268 androgen-induced and 1313 androgen-repressed genes, and significant overlap of these gene sets with LNCaP and VCaP cells [7]. PC3-AR cells were also analyzed by AR-ChIP, which revealed androgen-dependent AR binding to target genes [8]. We recently sequenced the LNCaP, VCaP, and PC3-AR cell lines to determine whether these prostate cancer models harbor deleterious mutations in the DNA repair machinery [7]. Here, we report and compare the features of the whole-genome sequence from the three cell lines.

## Materials and Methods

### Cell culture

LNCaP, VCaP, and PC-3 cells were purchased from ATCC. VCaP cells were grown in DMEM supplemented with 10% FBS and 1% antibiotic/antimycotic. LNCaP and PC-3 cells were grown in RPMI supplemented with 5% FBS and 1% antibiotic/antimycotic. PC3-AR cells, as described previously [7] were generated by stable lentiviral infection of full-length AR using a pWPI-GFP-FLAG-AR plasmid in which the GFP portion was swapped with the antibiotic selectable hygromycin resistance gene. All cells were incubated at 5% CO_2_ and 37 °C.

### Genome sequencing and alignment

For LNCaP, VCaP, and PC3-AR cell lines, genomic DNA was prepared using the Qiagen DNeasy kit. Libraries were prepared, and samples were sequenced by Hudson Alpha. DNA was aligned using BWA MEM [9] available in v0.7.17 to the b37+decoy reference sequence. The putative PCR duplicates were flagged using SAMBLASTER v0.6.9 [10], and a sorted BAM file was generated using SAMtools v1.9 [11].

### Variant calling

We used FreeBayes [12] to identify variants (single-nucleotide variants and small indels) in the three cell lines and filtered to keep all variants with a Phred-scaled variant quality greater than 20. We used vt [13] to decompose and normalize the variants, and VEP [14] to annotate the variants. BCFtools [15] was used to filter and summarize the resulting VCF file, and TAPES [16] was used to identify the pathogenic mutations in the variant calls.

### Determination of copy number states

HMMcopy [17] was used to assign read counts to 10 Kbp bins, and the copy number segments were identified using circular binary segmentation [18] of those bin counts.

### Pathway enrichment

We used SLAPEnrich [19] to identify the significantly mutated KEGG pathways in the cell lines.

### Structural variant detection

We used Meerkat [20] to identify the structural variants (SVs) in the three cell lines. In addition to detecting the structural variants, Meerkat uses sequence homology at the breakpoints to infer the mechanism that formed the SVs and has been applied to understand the landscape of SVs in several studies [20, 21]. To differentiate detected SVs from non-unique regions, we required SVs to have support from both discordant read-pairs and split-reads, and we limited SVs to the ones with <= 40 bps of homology around the breakpoint.

### Chromothripsis

We used ShatterSeek [21] to determine if the cell lines exhibit known signatures of chromothripsis.

## Results

### Copy number profiles and variant calls

Spectral karyotyping (SKY) of CaP cell lines has demonstrated aneuploid karyotypes with many chromosomal aberrations, including complex chromosomal rearrangements and a high degree of karyotype instability [22]. Figure 1A, B, C shows the distribution of read counts assigned to mappable bins of 10 Kbps for VCaP, PC3-AR, and LNCaP cells. The majority of the bins in VCaP cells can be assigned a copy number state of three, which agrees with previous reports that VCaP cells are near triploid [23]. The distribution of read counts in the PC3-AR cells shows similar separable peaks corresponding to integral copy number states. Previous reports suggest that PC3 cells are near triploid, though based on the assignment of copy number states, these PC3-AR cells appear to be tetraploid [24]. In contrast to VCaP and PC3-AR cells, the distribution of read counts in LNCaP cells does not show discrete integral peaks corresponding to copy number states. This non-separability of copy number states is in line with reports that LNCaP cells harbor multiple clones with naturally occurring differences in androgen sensitivity caused by spontaneously arising changes [1].

**Figure 1:**
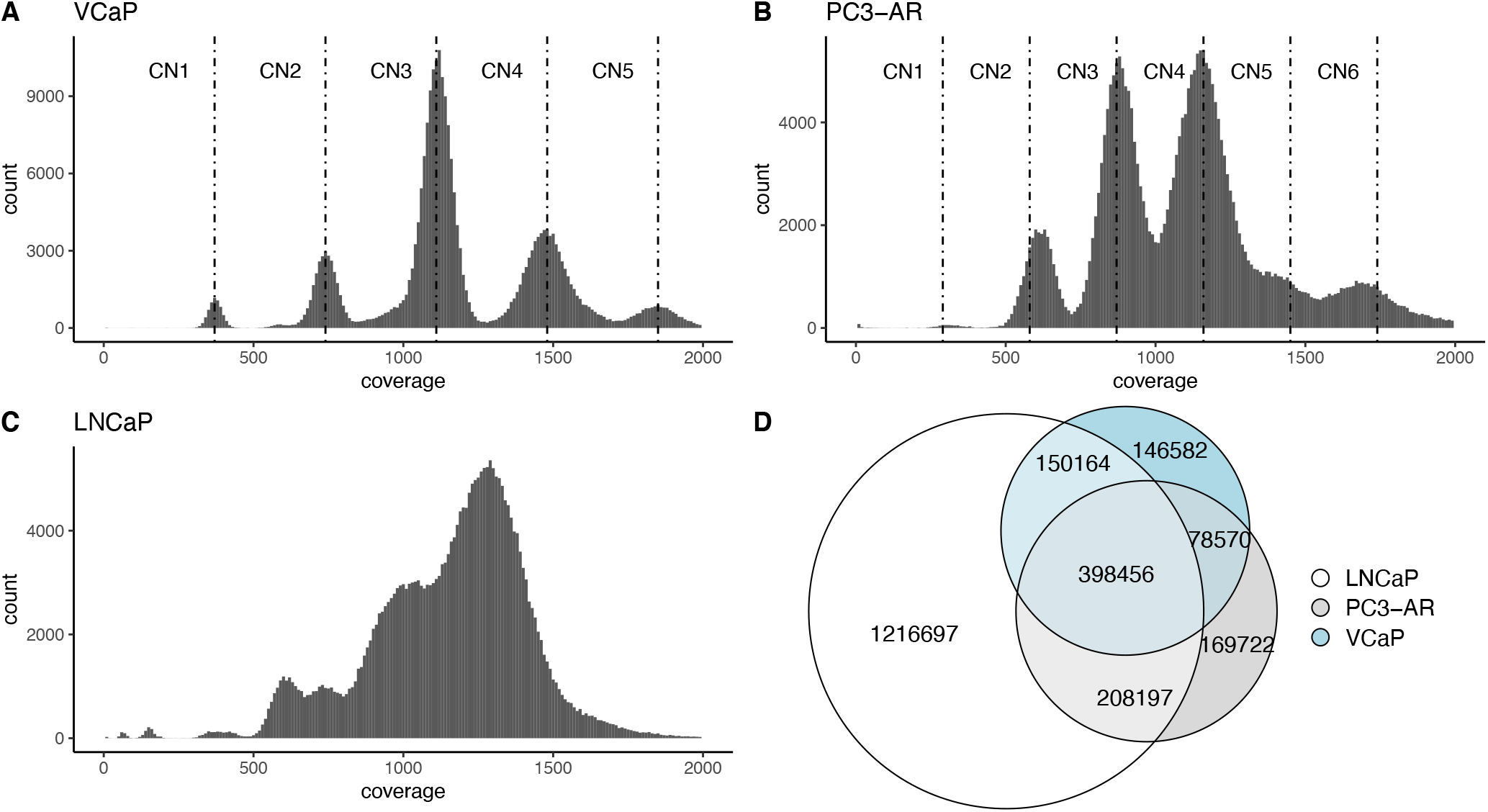
Distribution of read counts in mappable bins of 10 Kbps for (A) VCaP, (B) PC3-AR, and (C) LNCaP cells. VCaP and PC3-AR cells show integral copy number states. (D) Rare SNVs shared by the three cell lines.

Table 1 enumerates the protein-coding genes with zero copies in the three cell lines based on read count assignments in 10 Kbp bins. In LNCaP, we identified a ~300 Kbp homozygous deletion (chr2:47,677,007-47,986,422) that overlaps the region with the *KCNK12* gene and multiple exons of *MSH2*. Consistent with this observation, the LNCaP parental strain is known to lack *MSH2* expression. The *MSH2* protein dimerizes with *MSH6* to recognize base mismatches, and *MSH2* also dimerizes with *MSH3* to identify large insertion-deletion loops [25]. The deletion of *MSH2* in LNCaP likely affects the mismatch repair pathway in the cells. Among other genes, *USP10* and *PPP2R2A* are deleted in VCaP cells. The *USP10* gene controls the epigenetic changes induced by the androgen receptor (AR), and the lack of *USP10* protein is known to decrease the stability of AR targets and lead to poor outcomes in human CaP [26]. Similarly, *PPP2R2A* is hemizygously lost in ~42% of prostate adenocarcinomas, correlates with reduced expression, poorer prognosis, and an increased incidence of hemizygous loss (>75%) in metastatic disease [27].

**Table 1:** Genes with zero copies in LNCaP, VCaP, and PC3-AR cell lines.

| Cell line | Genes completely deleted in cell line |
| --- | --- |
| LNCaP | *MSH2, KCNK12* |
| VCaP | *PPP2R2A, BNIP3L, USP10, CRISPLD2, LCE3C, LCE3B* |
| PC3-AR | *CTNNA1, LRRTM2, SFTPA2, SFTPA1, MAT1A, DYDC1, DYDC2, PRXL2A, TSPAN14, PTEN, RNLS, LIPJ, LIPF, LIPK, RP11-473M10, KAT2A, CTD-2132N18.3, HSPB9, RAB5C, KCNH4, HCRT, GHDC, STAT5B, STAT5A, STAT3, CAVIN1, ATP6V0A1, OR4F17, PLPP2, MIER2, CIC, PAFAH1B3, LCE3C, LCE3B* |

Both PC3-AR and VCaP cells have deleted segments on chromosome 1 overlapping *LCE3B* and *LCE3C* genes. Even though these late cornified envelope genes are deleted in ~18.2% of prostate adenocarcinomas, their role in tumorigenesis is unclear [28]. The gene *PTEN* is entirely deleted in the PC3-AR cells. Deep deletion of *PTEN* is a known driver of CaP observed in ~40% of metastatic prostate adenocarcinoma patients and is associated with reduced expression of AR-regulated luminal epithelial genes [29]. *Capicua*, a known tumor suppressor and an evolutionarily conserved high-mobility group-box transcription factor, is also deleted in PC3-AR cells [30, 31]. The gene *CTNNA1*, known to be truncated in patients with hereditary diffuse gastric cancer (HDGC), is another gene with a known oncogenic role deleted in PC3-AR cells.

We next used FreeBayes [12] to identify variants (positions with SNVs and small indels) in the three cell lines. FreeBayes identified 4,268,194 variants in PC3-AR, 4,433,670 variants in VCaP cells, and 5,699,585 variants in LNCaP cells. A significant fraction of these mutations are likely from the germline of the patients from whom these cell lines were derived. Since tumors are typically enriched for rare and deleterious mutations, we filtered the calls to remove the mutations with an allele frequency greater than 0.01 in human populations in an attempt to identify putative somatic mutations. This left us with 854,945 variants in PC3-AR, 773,772 variants in VCaP and 1,973,514 variants in LNCaP cells. Figure 1D shows the overlap of these mutations in the three cell lines, again highlighting that LNCaP cells have a high somatic burden due to the heterogeneity of the cells.

### Pathogenic coding SNVs

We used TAPES [16] to identify pathogenic single-nucleotide variants (SNVs) in the protein-coding regions of the three cell lines using the criterion as defined by the American College of Medical Genetics (ACMG) [32]. We identified nine variants in the VCaP cell line, 17 variants in the PC3-AR cell line, and 351 pathogenic variants affecting 327 genes in the LNCaP cell line annotated as “Pathogenic” or “Likely Pathogenic” (Table S1).

Comparing the list of pathogenic variants (Figure 2A) reveals that LNCaP and PC3-AR both harbor such variants in *TP53*. This known tumor suppressor plays a pivotal role in genomic stability, cell cycle arrest, and other critical signaling pathways. 18% of patients diagnosed with prostate adenocarcinoma have an alteration in *TP53*. PC3-AR cells are hemizygous for chromosome 17p, and the only *TP53* allele has a frameshift deletion leading to premature termination of the protein product (*TP53:NM_001126112:exon5:c.414delC:p.K139Rfs*31*). Consistent with this result, PC3-AR does not express *TP53*. LNCaP cells, on the other hand, harbor a missense mutation in *TP53* (*TP53:NM_001126112:exon7:c.T700C:p.Y234H*) that overlaps a known somatic locus in multiple carcinomas. Even though TAPES did not annotate any mutations as pathogenic in the *TP53* gene of the VCaP cells, we identified a homozygous *c.742C>T (p.R248 W)* mutation that inactivates *TP53* and has been observed in prostate adenocarcinomas. That mutation is annotated as a variant of uncertain significance (VUS) by TAPES which assigns it a probability of 0.675 of being pathogenic.

**Figure 2:**
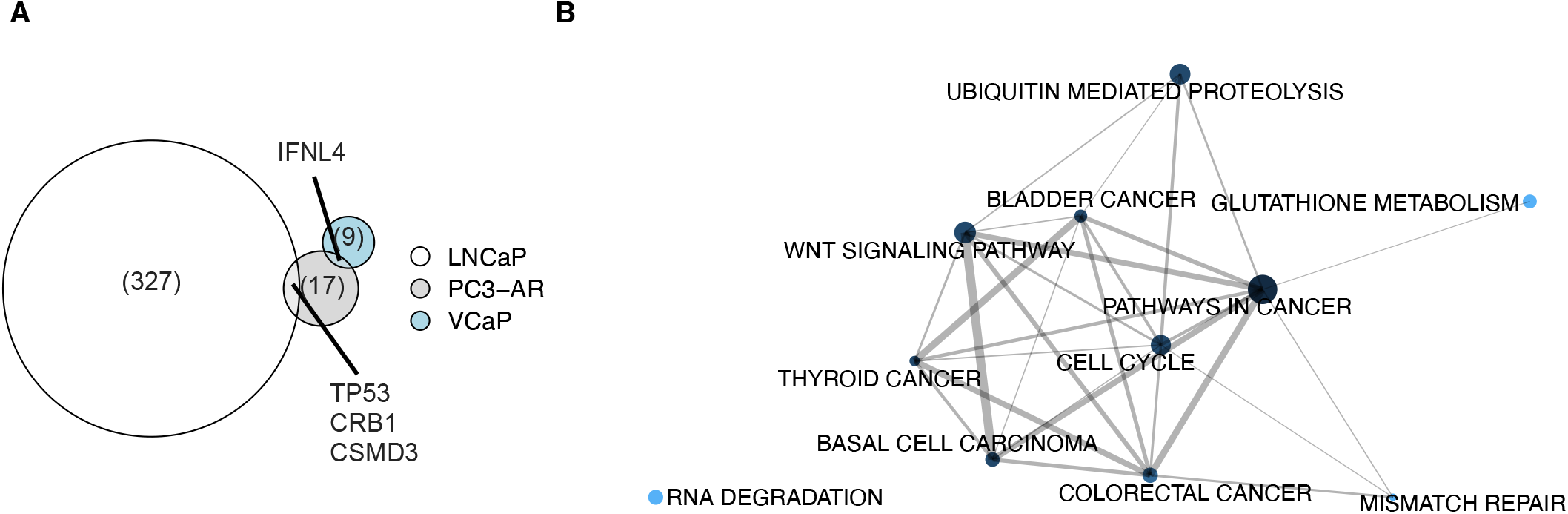
(A) Comparison of pathogenic variants identified in LNCaP, PC3-AR, and VCaP cells. (B) Pathways with recurrent pathogenic mutations. The size of the nodes is proportional to the number of genes in the pathway, and the thickness of the edges between two nodes reflects the overlap based on gene membership between them.

Besides *TP53*, LNCaP and PC3-AR also harbor pathogenic variants in *CRB1* and *CSMD3*, but these genes are not expressed in the prostate. *CRB1* encodes a protein that localizes to the inner segment of mammalian photoreceptors and controls the proper development of the polarity in the eye, and *CSMD3* is involved in dendrite development. VCaP and PC3-AR harbor a deltaG allele in the IFNL4 gene, which encodes for interferon IFN- λ4 protein and is associated with an impaired ability to clear the hepatitis-C virus [33].

We also found three pathogenic mutations in LNCaP cells that are annotated as relevant to human prostate cancers. We found a stop-gain in *CHEK2 (CHEK2:NM_001349956:exon5:c.G514T:p.E172X)*, a serine/threonine-protein kinase that is required for checkpoint mediated cell cycle arrest and activation of DNA repair and apoptosis in response to the presence of DNA double-strand breaks. We also observed a loss-of-heterozygosity (LOH) and a frameshift deletion in *PTEN*, leading to a stop-gain (*PTEN: NM_000314:exon1:c. 16_17del:p.K6Rfs*4*). *PTEN* is a known tumor suppressor that antagonizes the PI3K-AKT/PKB signaling pathway and is also entirely deleted in the PC3-AR cells. This gene is also homozygously deleted in the PC3-AR cells. Lastly, we found that LNCaP harbors a frameshift deletion in *RNASEL (RNASEL: NM_021133:exon2:c.471_474del:p.K158Rfs*6)*, which is associated with familial predisposition to prostate cancer [34].

Using SLAPenrich [19], we then identified the pathways that are recurrently modified in these cell lines. Figure 2B shows a network of overlapping KEGG pathways recurrently mutated in these cell lines with an FDR < 0.05. The nodes represent the pathways, and the thickness of the edges reflects the overlap between the pathways based on gene membership. Besides the pathways annotated as being involved in cancers (pathways in cancer, colorectal cancer, bladder cancer, and thyroid cancer, and basal cell carcinoma), we observed pathogenic mutations in glutathione metabolism, cell cycle, Wnt signaling pathway, and RNA degradation pathway (*HSPA9, PAPOLG*). We also found recurrent dysregulation of the mismatch repair pathway in the three cell lines. In the VCaP cell line, we identified a heterozygous frameshift insertion in the *MSH6* gene. In the LNCaP cell line, we found a homozygous deletion of exons 9-16 of *MSH2*, which results in a truncated protein, and a frameshift deletion in the *MSH3* gene. *MSH3* heterodimerizes with *MSH2* to form MutS-beta, which binds to the DNA mismatches and recognizes large insertion-deletion loops, whereas *MSH6* heterodimerizes with *MSH2* to form MutS-alpha, which binds to DNA mismatches and initiates repair. This analysis further demonstrates that even though the cell lines have accumulated independent passenger mutations, the pathogenic mutations are enriched for dysregulation of pathways with a known role in oncogenic processes.

### Structural variants

We used Meerkat [20] to identify the structural variants (SVs) in the three cell lines. The detected SVs include the germline SVs and the changes acquired by the cells during tumorigenesis and in vitro culture. In addition to identifying the structural variants, Meerkat uses sequence homology at the breakpoints to infer the mechanism that formed the SVs and has been applied to characterize the landscape of SVs in several studies. Table 2 shows the counts of the various types of SVs in each sample after filtering. VCaP cells have the highest number of structural variants among these cell lines, whereas LNCaP cells have the smallest.

**Table 2:** Count of various types of structural variants identified in LNCaP, PC3-AR, and VCaP cells.

| SV type | LNCaP | PC3-AR | VCaP |
| --- | --- | --- | --- |
| Deletion | 778 | 747 | 690 |
| Deletion with insertion | 151 | 223 | 244 |
| Deletion with inversion | 4 | 14 | 34 |
| Insertion | 30 | 28 | 29 |
| Interchromosomal translocation | 202 | 211 | 223 |
| Inversion | 88 | 159 | 276 |
| Tandem duplication | 97 | 176 | 141 |

Deletions are usually the result of DNA double-strand break repair. Transposable element insertion (TEI) is the dominant mechanism of deletion-like SV formation in the germline, whereas non-homologous end joining (NHEJ) and alternative end-joining (alt-EJ) have been reported as the dominant mechanisms in somatic cells [20]. We used Meerkat to predict the mechanism of formation for deletion-like fragment joints in these cells. Figure 3A shows the count of deletion-like joins in the three cell lines predicted to be created via various mechanisms, and Figure 3B shows the size distribution of these deletion-like joins. Based on these results, deletions generated via NHEJ and fork stalling and template switching/microhomology-mediated break-induced repair (FoSTeS) tend to be longer than those generated via TEI, which probably reflect germline variants. Looking at interchromosomal translocations and inversion-like joins in the cell lines, we can see that most breakpoints lack homology and are likely created via NHEJ/alt-EJ/FoSTeS mechanisms that are the dominant repair mechanisms in tumors.

**Figure 3:**
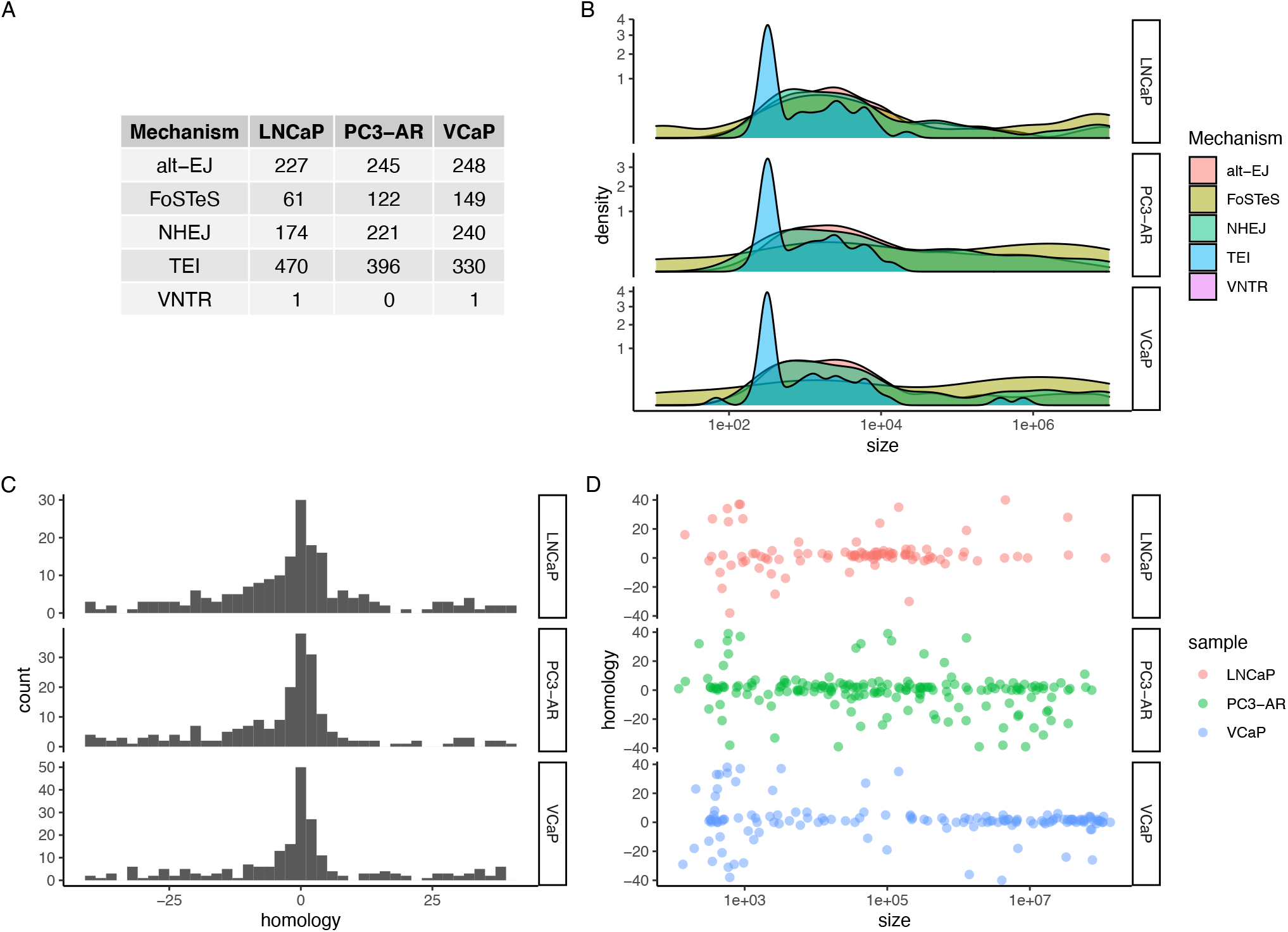
Characteristics of SVs in LNCaP, VCaP, and PC3-AR cells. A) Count of deletion generated via various mechanisms. (B) Size distribution of deletions stratified by the putative mechanism that generated them. (C) Homology around breakpoints for Interchromosomal translocations detected in three cell lines. A negative homology indicates the insertion of bases. (D) Size and homology for inversion-like structural variants.

The *TMPRSS2-ERG* gene fusion is the most frequent genomic alteration in prostate cancer, resulting in overexpression of the transcription factor *ERG*. Several studies have reported that VCaP cells harbor that fusion, but the landscape of putative gene fusions in these commonly used CaP cell lines has not been reported. We identified 533 gene fusions in the VCaP cell line, 553 gene fusions in the PC3-AR cell line, and 476 gene fusions in the LNCaP cell line (Table S2). As expected, most of the gene fusions do not change the protein products (Table 3). In the VCaP cells, *TMPRSS2* is involved in several rearrangements, including two intragenic rearrangements and two intergenic rearrangements with *ERG* and *C16orf7*.Based on the micro-homology around the breakpoints, it appears that the *TMPRSS2-ERG* fusions are generated via fork stalling and template switching mechanisms. We also observed a *ZBTB20-LSAMP* gene fusion in the VCaP cell line, arising from a duplication that leads to a UTR swap.

**Table 3:** Gene-gene fusions in LNCaP, VCaP, and PC3-AR cells.

| Fusion type | LNCaP | PC3-AR | VCaP |
| --- | --- | --- | --- |
| head-head/tail-tail | 70 | 109 | 120 |
| In frame | 16 | 24 | 13 |
| No impact | 336 | 322 | 315 |
| Out of frame | 20 | 38 | 29 |
| Unknown | 6 | 9 | 9 |
| UTR swap | 28 | 51 | 47 |

In the LNCaP cells, we found evidence for a *BLNK-TLL2* gene fusion that has been reported in cervical cancer and a *BMPR2-FAM117B* in-frame gene fusion that has been reported in breast adenocarcinoma [35]. In the PC3-AR cells, we found an *IMMPL2L-DOCK4* gene fusion that has been reported in prostate adenocarcinoma (PRAD), stomach adenocarcinoma (STAD), and esophageal carcinoma (ESCA). We also identified a *KDM5B* intragenic rearrangement, which has been reported in colonic neoplasms, and a *TFCP2-SMAGP* gene fusion reported in lung squamous cell carcinoma (LUSC). We also detected repeated deletions in the *PTPRD* genes, all of which are predicted to have been generated via alt-EJ and NHEJ mechanisms.

### Chromothripsis

Chromothripsis involves the shattering of one or a few chromosomes followed by fragment reassembly. It is defined based on (1) occurrence of genomic rearrangements in localized chromosomal regions, (2) copy number changes alternating a few copy number states, and (3) the alternation between regions where heterozygosity is preserved with regions displaying loss of heterozygosity. Chromothripsis has previously been reported in the VCaP cells on the q arm of chromosome 5 [23]. A recent study analyzed the breakpoints involved in canonical chromothripsis events with interspersed LOH and concluded that NHEJ has a principal role in DNA repair, with partial contributions from MMBIR/FoSTeS or alt-EJ [21].

We used ShatterSeek [21] to detect the presence of chromothripsis in the three cell lines. We did not find any evidence of chromothripsis in LNCaP cells. For VCaP cells, ShatterSeek reported high confidence of chromothripsis in chromosome 5, chromosome 9, chromosome 13, chromosome 14, chromosome 16. Figure 4A shows the fragment joins and copy numbers observed in chromosome 5 of VCaP cells. The affected region on 5q has 358 intrachromosomal structural variants in 182 segments, with at least a contiguous oscillation between two copy number states for 12 of those segments. Of the 170 deletion-like fragment joins in the region, we find that 85 exhibit signatures consistent with being generated via fork stalling and template switching mechanisms (FoSTeS), whereas the majority of the remaining breakpoints were split between NHEJ (35) and alt-EJ (36) mechanisms. As reported by Alves et al. [23], we did not find any evidence of positive selection of in-frame fusion transcripts in this region.

**Figure 4:**
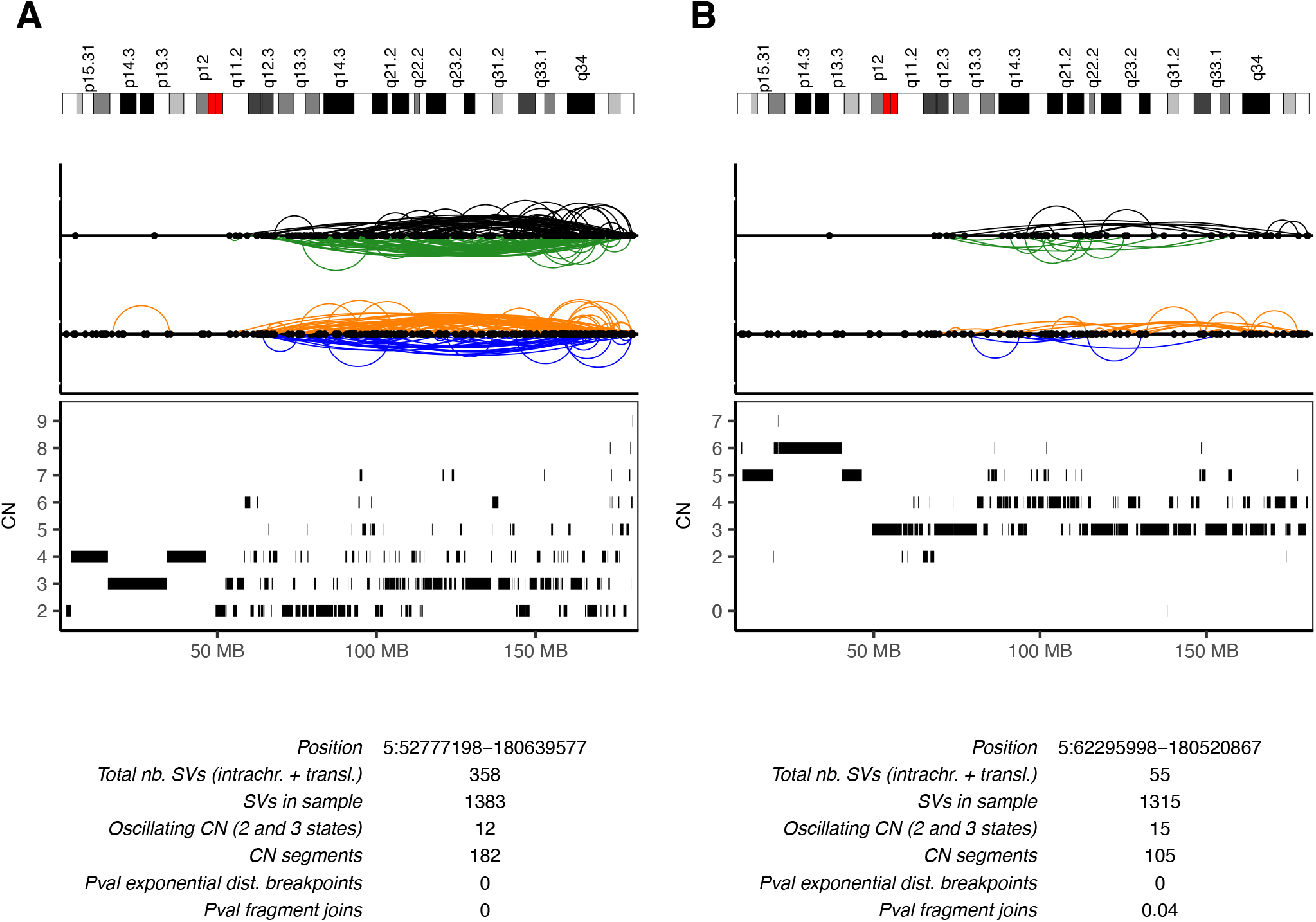
Chromothripsis of chromosome 5, arm q is observed in both VCaP and PC3-AR cells.

Surprisingly, we also found evidence of chromothripsis in chromosome 5 and chromosome 8 of the PC3-AR cells. Figure 4B shows the fragment joins, copy number states for chromosome 5 of the PC3-AR cells. The affected region on 5q has 55 intrachromosomal structural variants in 105 segments, with at least one contiguous oscillation between two copy number states for 15 of those segments. Of the 57 deletion-like fragment joins in the region, we identified that 25 are generated via fork stalling and template switching mechanisms (FoSTeS), 11 are generated via alt-EJ, and six are generated via NHEJ.

## Discussion

Whole-genome ploidy and modal chromosome number have been characterized for most cancer cell lines. Using mappable bins of 10 Kbps, we found that most bins in VCaP and PC3-AR can be assigned to integral copy number states. However, the same is not valid for LNCaP cells, which appear to consist of multiple cellular subpopulations. Other studies have reported that LNCaP cells harbor multiple clones with naturally occurring differences in androgen sensitivity caused by spontaneously arising changes. This instability and molecular heterogeneity in LNCaP cells could explain how multiple strains have been developed from the LNCaP parental strain, including cells under selection for growth under conditions of androgen depletion. As an example, one of LNCaP derivative cell lines (LNCP-Abl) differentially displays a resistance to enzalutamide [37].

Our study used TAPES [12] to identify the pathogenic coding mutations in the three cell lines. Even though this highlighted several essential pathways, such as the mismatch repair pathways that are recurrently mutated in the cell lines, this analysis could still have missed several variants relevant to the disease. For example, the homozygous *c.742C>T (p.R248 W)* mutation in the VCaP cells was annotated as a variant of uncertain significance by TAPES, even though it has been shown that the mutation inactivates *TP53*. This result underscores the challenges of variant prioritization and highlights the difficulties in determining somatic variants in a tumor-only sample.

In this report, we identified several structural variants in these cell lines. Among them was a *ZBTB20-LSAMP* gene fusion in the VCaP cell line, arising from a duplication that leads to a UTR swap. In another study, *LSAMP* locus rearrangements were found to be associated with African American ethnicity, and *LSAMP* deletion was found to be correlated with rapid disease progression [38]. Our analysis suggests that rearrangements of *LSAMP* locus might not be specific to African Americans.

Deletion of chromosome 5q is common in prostate cancer, affecting between 6% to 19% of prostate cancers, and is linked to aggressive course of disease [39]. We found evidence of chromothripsis in chr5q in both PC3-AR and VCaP cells. About half of the breakpoints exhibit evidence of generation via fork stalling and template switching mechanisms (FoSTeS). This contrasts with an analysis of various tumors, which concluded that more than half of the breakpoints appear to be generated via NHEJ or alt-EJ. We did not observe any complete gene loss in the regions affected by chromothripsis. Since VCaP and PC3-AR are polyploid cell lines, this suggests that the shattering event occurred after the duplication events.

We also found inactivating *TP53* mutations in both PC3-AR and VCaP cells. *TP53* malfunction has been reported as a predisposing factor for chromothripsis [21]. LNCaP cells also have a pathogenic missense mutation in *TP53*. Still, they do not show any evidence of chromothripsis, suggesting that *TP53* deficiency rather than malfunction has a role in predisposing cells to chromothripsis. It also remains to be determined whether chromosome-specific properties or the non-random localization of chromosomes play a role in the preferential target of specific chromosomes by chromothripsis.

## Author contributions

CY, BMP, and AR designed the research; KS and AR conducted computational analyses; BMP provided funding acquisition; BMP and AR wrote the manuscript. All authors read and approved the final manuscript.

## Competing interest statement

The authors declare that they have no competing interests.

## Notes

### Competing Interest Statement

The authors have declared no competing interest.

